# Stingless bee floral visitation in the global tropics and subtropics

**DOI:** 10.1101/2021.04.26.440550

**Authors:** Francisco Garcia Bulle Bueno, Liam Kendall, Denise Araujo Alves, Manuel Lequerica Tamara, Tim Heard, Tanya Latty, Rosalyn Gloag

## Abstract

Bees play a key role in maintaining healthy terrestrial ecosystems by pollinating plants. Stingless bees (Apidae: Meliponini) are a diverse clade of social bees (>500 species) with a pantropical distribution spanning South and Central America, Africa, India, Australia and Asia. They are garnering increasing attention as commercially-beneficial pollinators of some crops, yet their contribution to the pollination of native plants in the tropics and subtropics remains poorly understood. Here we conduct a global review of the plants visited by stingless bees. We compile a database of reported associations (flower visits) between stingless bees and plants, from studies that have made either direct observations of foraging bees or analysed the pollen stored in nests. Worldwide, we find stingless bees have been reported to visit the flowers of plants from at least 220 different families and 1465 genera, with frequently reported interactions for many of the tropic’s most species-diverse plant families including Fabaceae, Asteraceae, Rubiaceae, Malvaceae, Lamiaceae, Arecaceae, Euphorbiaceae, Poaceae, Apocinaceae, Bignoniaceae, Melastomataceae and Myrtaceae. The list of commonly-visited plant families was similar for the stingless bee fauna of each of three major biogeographic regions (Neotropical, Afrotropical and Indo-Malayan-Australasian), though we detected differences in the proportional use of plant families by the stingless bees of the Indo-Malayan-Australasian and Neotropical regions, likely reflecting differences in the available flora of those regions. Stingless bees in all regions visit a range of exotic species in their preferred plant families (crops, ornamental plants and weeds), in addition to native plants. Although most reports of floral visitation on wild plants do not confirm effective pollen transfer, it is likely that stingless bees make at least some contribution to pollination for the majority of plants they visit. In all, our database supports the view that stingless bees play an important role in the ecosystems of the global tropics and subtropics as pollinators of an exceptionally large and diverse number of plants. This database also highlights important gaps in our knowledge of stingless bee resource use that may help focus future research efforts.

## Introduction

Animal pollination is critical to ecosystem functioning and service provisioning in terrestrial ecosystems globally (Fontaine et al., 2005; Klein et al., 2006; Potts et al., 2016). While a diverse range of vertebrates and arthropods may pollinate plants, the majority of plant species (65-80%) rely on insects as their primary pollinators (Buchmann et al., 2012; Ollerton et al., 2011). As such, ongoing declines in insect pollinator populations in recent decades have the potential to cause synergistic declines in their pollinator-dependent plants (Biesmeijer et al., 2006; Potts et al., 2016).

The plants of the species-rich tropics and subtropics may be particularly vulnerable to changes in insect populations, as the flora of these regions are more dependent on pollinators than those of temperate-zones (Ollerton et al., 2011). In addition, insects in the tropics are predicted to be the least capable of rapidly adapting to changing climates, and thus the most at risk of extinction (Kellermann et al., 2012). In order to mitigate the impacts of pollinator declines in the tropics and subtropics, we first require a greater understanding of pollinator ecology in these regions (Vanbergen et al., 2013), and the relationship between floral resources and pollinators at landscape scales (Kleijn et al., 2008; Vanbergen et al., 2013).

The stingless bees (Hymenoptera: Apoidea: Anthophila: Meliponini) are social corbiculate bees native to the world’s tropical and subtropical regions (Michener, 1979, 2000; Roubik, 1992b). Like other social bees (*e.g*., honey bees, *Apis* sp.), stingless bees are abundant flower visitors in the ecosystems they inhabit, because each colony contains thousands to tens of thousands of workers (Hepburn et al., 2011; Michener, 1974; Nogueira Neto, 1997). Worldwide, there are an estimated 516 species of stingless bees in 60 genera (Rasmussen et al., 2007, 2010; Schuh, 2010). These are divided between three monophyletic clades that diverged from each other 50-70 millions years ago and whose distributions today largely correspond to three major biogeographic regions: Neotropical, African and Indo-Malayan-Australasian (Rasmussen et al., 2010). The Neotropics harbours the greatest stingless bee diversity, with 417 known species (*c*. 80% total diversity; (Moure, 2012).

The high global species diversity of stingless bees is also reflected in their morphological and behavioural diversity (Michener, 2000). They range in body size from just 2mm (some *Trigonisca* spp.) (Roubik, 2018) to 15 mm (*Melipona fuliginosa;* (Camargo et al., 2008). They may nest in tree cavities, termite nests, or underground (Grüter, 2020). In the absence of an effective sting, they have evolved varied defence mechanisms including acid discharge (Roubik, 2006), suicidal biting (Shackleton et al., 2015) and sticking resin (Lehmberg et al., 2008). And they show a variety of social habits, whereby colonies may have single queens (monogynous; most species) or multiple queens (facultatively polygynous; *Melipona bicolor*) (Velthuis et al., 2006; Vollet-Neto et al., 2018), workers may lay eggs regularly or be completely sterile (Bueno et al., 2020; Sommeijer et al., 1999; Tóth et al., 2004), and the workers of some species include a “solider caste” (Grüter et al., 2012). All stingless bees, however, share a need to visit flowers for nectar to feed themselves, and almost all also collect pollen to provision their offspring (excluding a handful of Neotropical species that feed their offspring carrion) (Mateus et al., 2004; Sakagami et al., 1993). Stingless bees therefore contribute to pollinating the native flora of forests throughout the tropics and subtropics (Roubik, 1992a).

Most research of stingless bee floral visitation to date has focused on their role as pollinators of crops (Giannini et al., 2015; Ish-Am et al., 1999; Nates Parra, 2016). Similar to honey bees, stingless bees can be readily kept and transported in hives (Nogueira Neto, 1997; Wille, 1983) meaning they can be introduced to orchards when in flower and then reallocated (Giannini et al., 2020). They are effective pollinators of a variety of tropical and subtropical crops species, including *açaí* palm (*Euterpe oleracea*), coconut (*Cocos nucifera*), coffee (*Coffea arabica*), macadamia (*Macadamia* spp.) and rambutan (*Nephelium lappaceum*) (Giannini et al., 2015; Heard, 1999; Slaa et al., 2006). They are also adapted to the local conditions in many regions where these crops are grown (Jaffé et al., 2015), with wild stingless bees providing valuable free pollination services. For example, in Australia, wild stingless bees (*Tetragonula carbonaria*) are as effective as managed honey bees at pollinating blueberry crops (*Vaccinium* spp.) (Kendall et al., 2020). In most parts of the world, however, our understanding of the relationship between stingless bees and non-crop plants is comparatively incomplete (Campbell et al., 2019; Roubik, 1995). Stingless bees actively forage on diverse floral resources throughout the year (Kleinert et al., 2012; Roubik, 1982, 1992a), but are proposed to have stronger interactions with some groups of native plants than others (Grüter, 2020). Indeed, at least some genera (*e.g., Melipona* spp. in the Neotropics) show a clear preference for some floral resources over others, irrespective of availability (Antonini et al., 2006; Kleinert et al., 2012; Ramalho et al., 1991; Ramalho et al., 1990; Vossler, 2013). Such preferences could arise because they confer a benefit to stingless bees via a reduction in interspecific competition (*e.g*., they focus on resources neglected by *Apis* spp.), or they may simply select those flowers that produce nectar and pollen in most abundance (Antonini et al., 2006; Ramalho et al., 1989a).

Here, we aim to consolidate current knowledge of stingless bee floral visitation of plants at the regional and global scale, by creating a database of reported floral visitation by stingless bees. From this database, we assess: (i) the diversity of plant families and genera used as food sources by stingless bees in each of three major biogeographical regions: Neotropics, Afrotropical and Indo-Malayan-Australasian; (ii) the most frequently-used plant families (according to number of genera visited), as a proxy for the broad floral preferences of stingless bees; and (iii) the endemism to a particular region, native or exotic status, and growth type of plants commonly used as forage by stingless bees. We also provide a reference list of those plants for which stingless bees have been experimentally confirmed to be pollinators (both crops and wild plants). Our database is intended as a first step towards a richer understanding of how stingless bees contribute to ecosystem functioning in the tropics and subtropics, and provides an online resource for further studies on plant-pollinator interactions.

## Method

### Database of floral visitation by stingless bees

To build a database of reported interactions between stingless bees and flowering plants, we conducted a search of peer review journals, books, student theses and conference abstracts. We used the keywords Meliponini, pollination, floral preferences and stingless bees to search Scopus, Web of Science and Google Scholar (first 200 references per search; May 2020). We likewise searched unpublished literature in Spanish and Portuguese from library databases and the University repositories of The National University of Colombia, The University of Costa Rica and Coordination for the Improvement of Higher Education Personnel (CAPES) of Brazil (Catálogo de Teses e Dissertações, from 2013 to now;(CAPES, 2016), plus the Conference abstracts from the annual Brazilian Bee Meeting “Encontro sobre Abelhas” (1994 to 2018). Finally, we included reported interactions from Brazil’s online index of Bee-Plant Interactions (A.B.E.L.H.A., 2017), and from three books: ‘Pot Pollen’(Vit et al., 2018), ‘Atlas of Pollen and Plants used by Bees’ (da Silva et al., 2020) and ‘Pollination of cultivated plants in the tropics’ (Roubik, 1995). We only included literature that was available online.

From each source, we collated the following information for each stingless bee species-flower interaction: biogeographic region (Neotropical, Afrotropical, Indo-Malayan-Australasian) and country where the interaction was reported, plant species, genus and family, bee species and type of interaction (either floral visitation or pollination). Floral visitation included cases where the researcher directly observed bees visiting flowers, or where visitation was inferred from palynological study (*i.e*., pollen resources collected from colonies or off the legs of returning foragers). An interaction was scored as pollination only if the study confirmed the bee pollinated the plant. We did not consider pollination efficiency (*i.e*., single visit efficiency of pollen deposition, fruit set or seed set; (King et al., 2013)) as only a handful of studies in our database reported such detail. Finally, for the 15 plant families with the greatest number of genera visited by stingless bees, we retrieved the native distribution of each genera and the growth type using Plants of the World Online (POWO, 2019) and noted whether the genus was documented as introduced in the region of the reported interaction according to the Centre for Agriculture and Bioscience International (CABI, 2020).

We considered commonly used plant families to be those in which stingless bees were reported to visit the most genera within the family. For the Neotropical region, we also accessed data on the total number of genera per plant family known to occur in the region via Neotropikey (Milliken, 2009), an online key developed to identify and inform about the flowering plants in the Neotropical region. This allowed us to account for high richness of genera in some plant families, by considering the proportion of locally-occurring genera in a family that were visited by stingless bees.

To confirm our database used current scientific names of all stingless bees, we checked names against the “Catalogue of Bees (Hymenoptera, Apoidea) in the Neotropics” (Moure, 2012), the “Catalogue of Afrotropical bees (Hymenoptera: Apoidea: Apiformes)” (Eardley et al., 2010) and the “Catalogue of the Indo-Malayan/Australasian stingless bees (Hymenoptera: Apidae: Meliponini)” (Rasmussen, 2008). For plant species, we cross-checked names against those listed on the Missouri Botanical Garden’s Tropicos website (Missouri Botanical Garden, 2020), The Plant List (The Plant List, 2010), Plants of the World Online (POWO, 2019), the Global Biodiversity Information Facility (GBIF) and the R package *taxize* v0.9.98 (Chamberlain et al., 2013).

### Analyses

We visualised plant ~ pollinator networks for the top ten most abundant bee genera and plant families using chord diagrams (*circlize* package v0.4.1; (Gu et al., 2014)). We also provide example flower types for each plant family in chord diagrams, based on (Simpson, 2010), to show the diversity of flower types utilized by stingless bees.

We tested whether the plant families visited by stingless bees varied between biogeographical regions (Neotropical, Afrotropical and Indo-Malayan-Australasian) using a two-step approach. First, we transformed our databases to interaction matrices of bee genera (rows) ~ plant family (columns, count of the number of visited genera) and then calculated the Bray-Curtis dissimilarity between bee genera using the *vegan* package v2.5-6 (Oksanen et al., 2013). Second, we compared compositional differences in plant use (at family level) between biogeographical regions using a pairwise PERMANOVA (Anderson, 2001; Martinez Arbizu, 2017). We clustered the bee genera from each of the three regions and compared the composition of reported plant families visited between each of the regions. We adjusted *P*-values using the false discovery rate (FDR) method to account for multiple comparisons (Benjamini et al., 1995). We visualised differences in the interactions between stingless bees and plant families in two-dimensional space with non-metric multidimensional scaling (nMDS) ordination. We conducted all data analyses in R v4.0.2 (Core Team, 2013).

## Results

Our database includes 19,768 bee-flower interactions reported in 597 studies (**Table S1**). In all, 53% (286/538) of species of stingless bees were represented by at least one reported interaction in the database; this spanned 53% (222/417) of Neotropical stingless bee species, 68% (22/32) of Afrotropical species and 47% (42/89) of Indo-Malayan-Australasian species. The great majority of reported interactions were floral visitation records (19,004 interactions, 96%), with the remainder (764 interactions, 4%) confirming that bee visitation resulted in pollination.

The majority of reviewed studies were carried out in the Neotropics (16899 interactions reported in 516 studies; 85% of all interactions), and particularly Brazil (13617 interactions); **Figure 1**. This reflects in part the higher diversity of stingless bees in this region (*e.g*., half of the world’s stingless bee genera are found in Brazil), and in part the intensity of research to date on the Neotropical stingless bee fauna, relative to that of the Afrotropical (1068 interactions reported in 17 studies; 5% of database interactions) and Indo-Malayan-Australasian regions (1801 interactions reported in 63 studies; 10% database interactions); **Figure 1**. Although our database included some “grey literature” (conference abstracts and student theses) from the Neotropics but not from other regions, the proportion of total interactions reported from the Neotropics was similarly high even if we included only bee visitations that were published in international journals.

**Figure 1.**
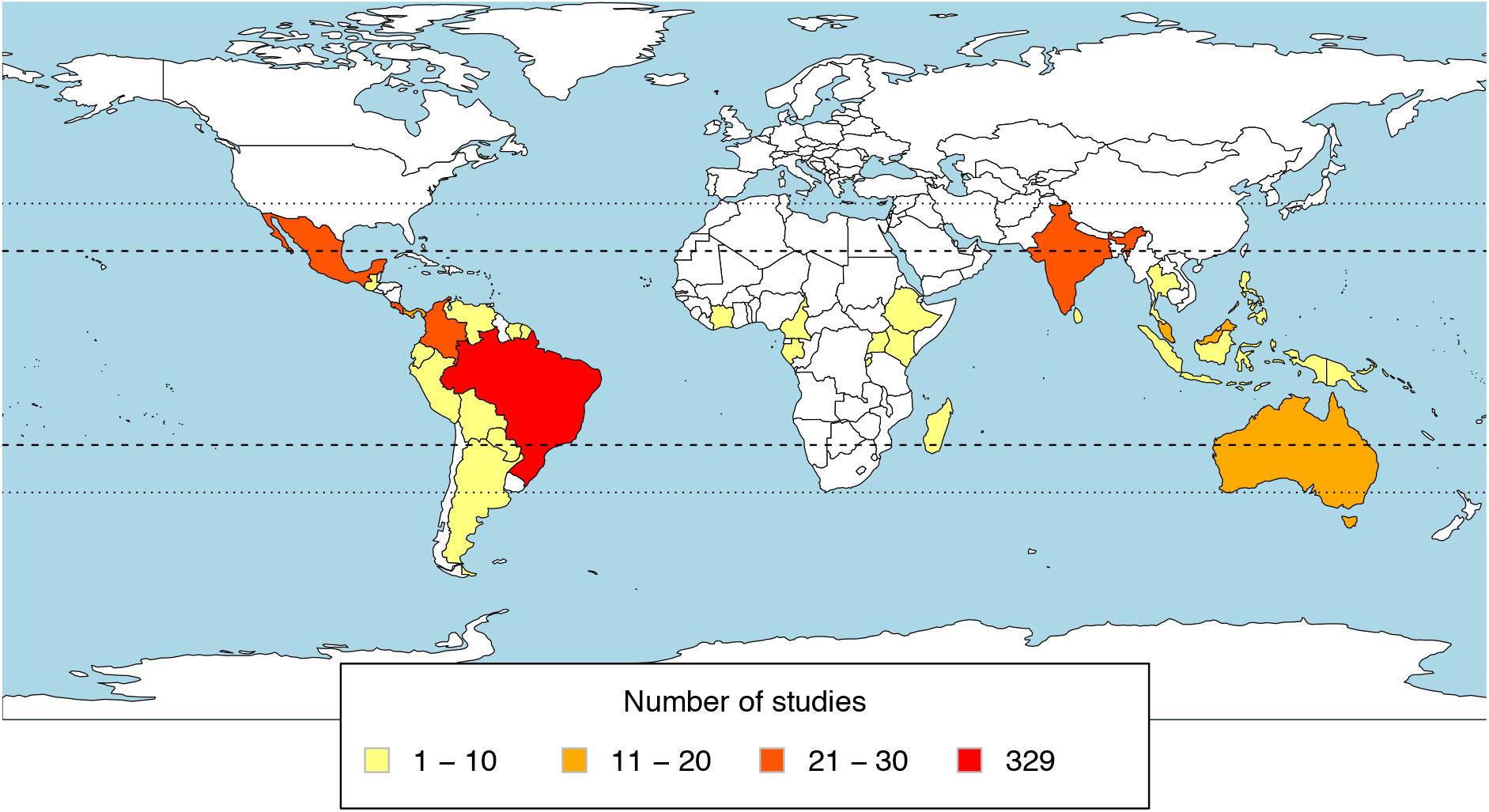
To visualise bias in reported stingless bee-plant interactions in the available literature, we plotted the number of studies that recorded interactions between stingless bees and flowering plants at a country level per region included in our database using *rworldmap* (v.1.3-6, South 2011). Dashed and dotted lines indicate the latitudinal range of the Tropics and Subtropics respectively.

### Plant genera and families visited by stingless bees

Stingless bees worldwide are reported to forage from the flowers of 1465 genera of plants in 220 families worldwide (52% of all angiosperm families); 200 families in the Neotropics, 77 in the Afrotropics and 132 in the Indo-Malayan-Australasian region (Sup Material **Table S1)**. Individual stingless bee species were reported to forage on 31 ± 60 (SD) plant genera (range: 1 – 532, with the maximum value reported for the Neotropical species *Trigona spinipes*).

The ten plant families with the largest number of genera visited by stingless bees were Fabaceae (legumes; N=161), Asteraceae (daisies; N=134), Rubiaceae (madders; N=65), Malvaceae (mallows; N=56), Lamiaceae (mints; N=43), Arecaceae (palms; N=41), Euphorbiaceae (spurges; N=41), Poaceae (grasses; N=34), Apocynaceae (dogbanes), Bignoniaceae (bignonias), Melastomataceae (melastomes), Myrtaceae (myrtles) (the last four, N=27 each); **Figures 2A-C, Figure 3, (**Sup material **Table S2, Fig. S1)**. All these families are highly diverse in number of genera and species, and all have pantropical distributions (Bramley et al., 2014).

**Figure 2A.**
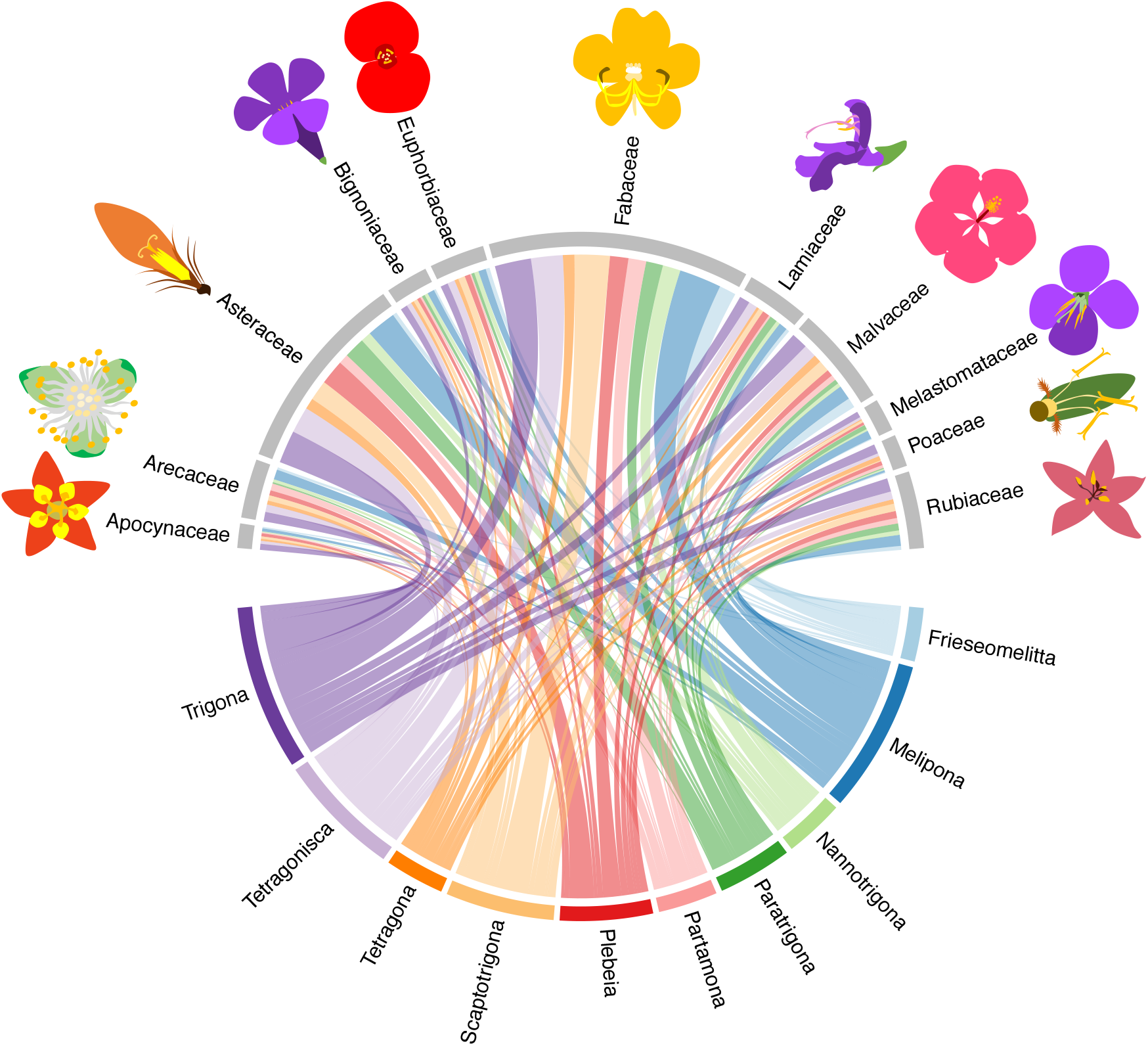
A visitation network of stingless bees to flowering plants in the Neotropical region, showing the 10 stingless bee genera and 10 plant families with the most reported interactions in the literature. Example flower types for each plant family are shown. Details of all interactions are given in Table S6 (Supplementary material).

**Figure 2B.**
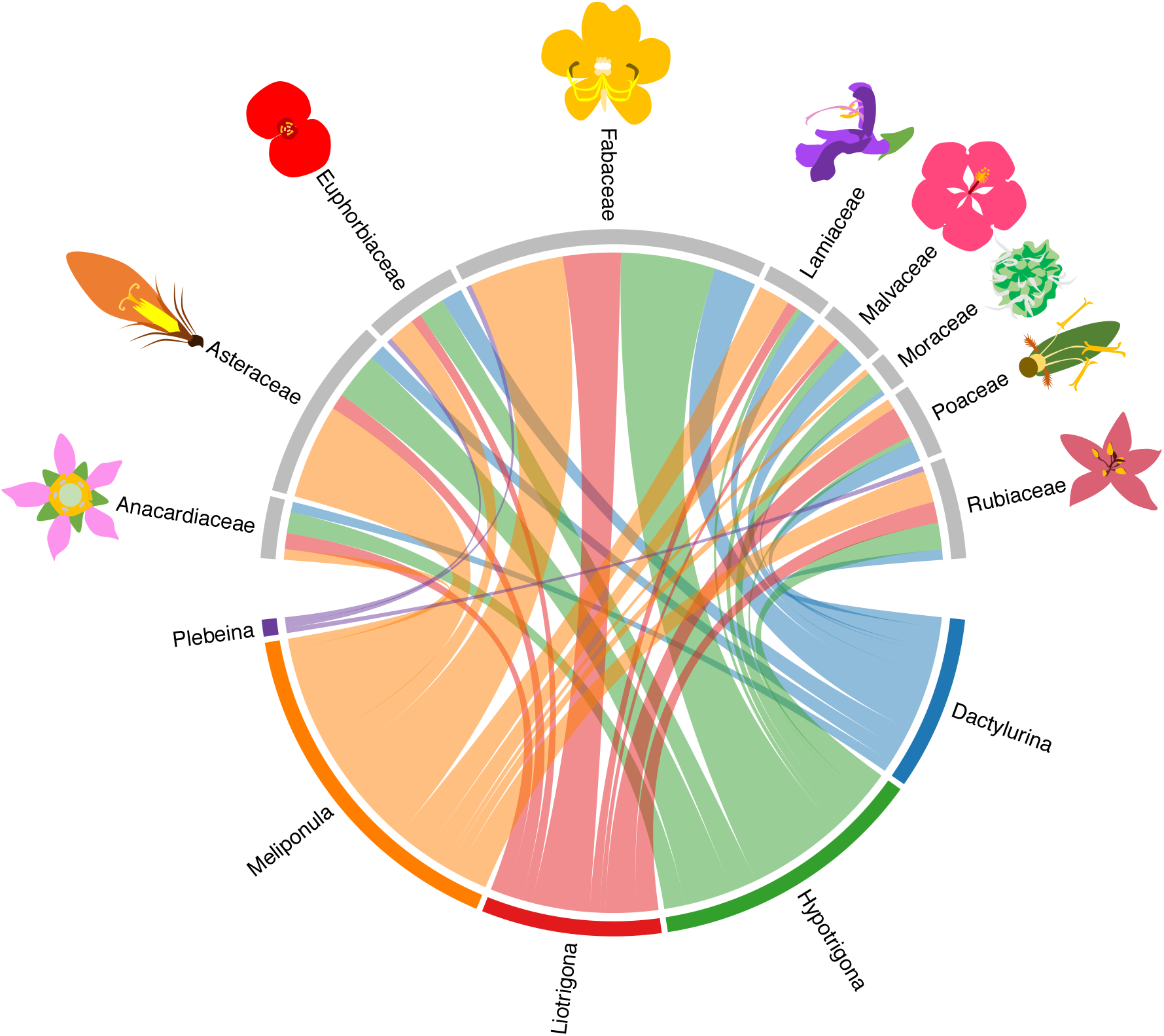
A visitation network of stingless bees to flowering plants in the Afrotropical region, showing the 5 stingless bee genera and 10 plant families with the most reported interactions in the literature. Only 9 families are shown here, otherwise to cramped, because the 10 had equally 5 families with X genera (Amaranthaceae, Arecaeae, Rhamnaceae, Rutaceae, Sapindaceae) Example flower types for each plant family are shown. Details of all interactions are given in Table S6 (Supplementary material).

**Figure 2C.**
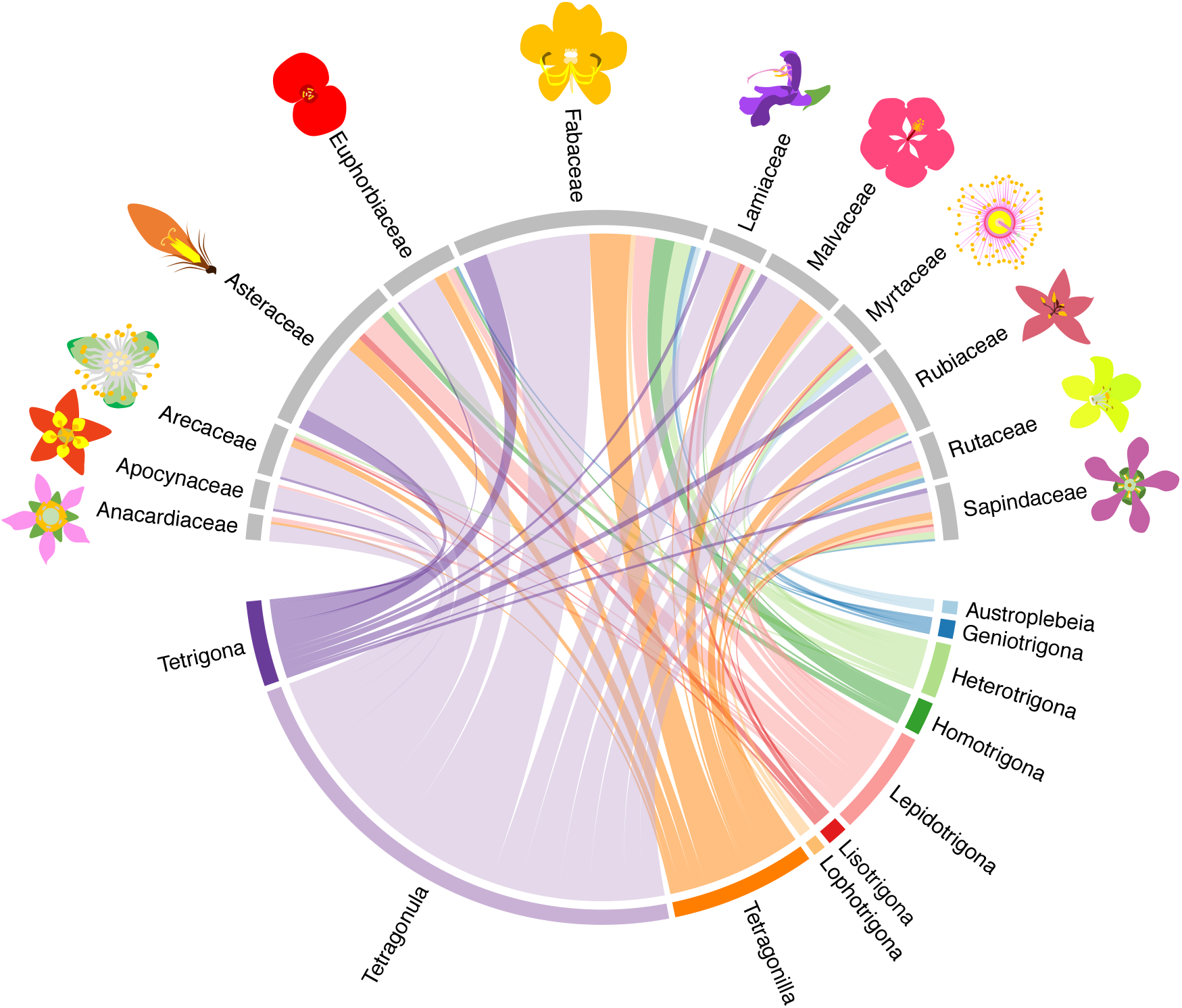
A visitation network of stingless bees to flowering plants in the Indo-Malayan-Australasian region, showing the 10 stingless bee genera (bottom; each represented by a different colour) and 10 plant families (top) with the most reported interactions in the literature. Bars connecting bee genera and plant families indicate the reported interactions. Example flower types for each plant family are shown. Details of all interactions are given in Table S6 (Supplementary material).

**Figure 3.**
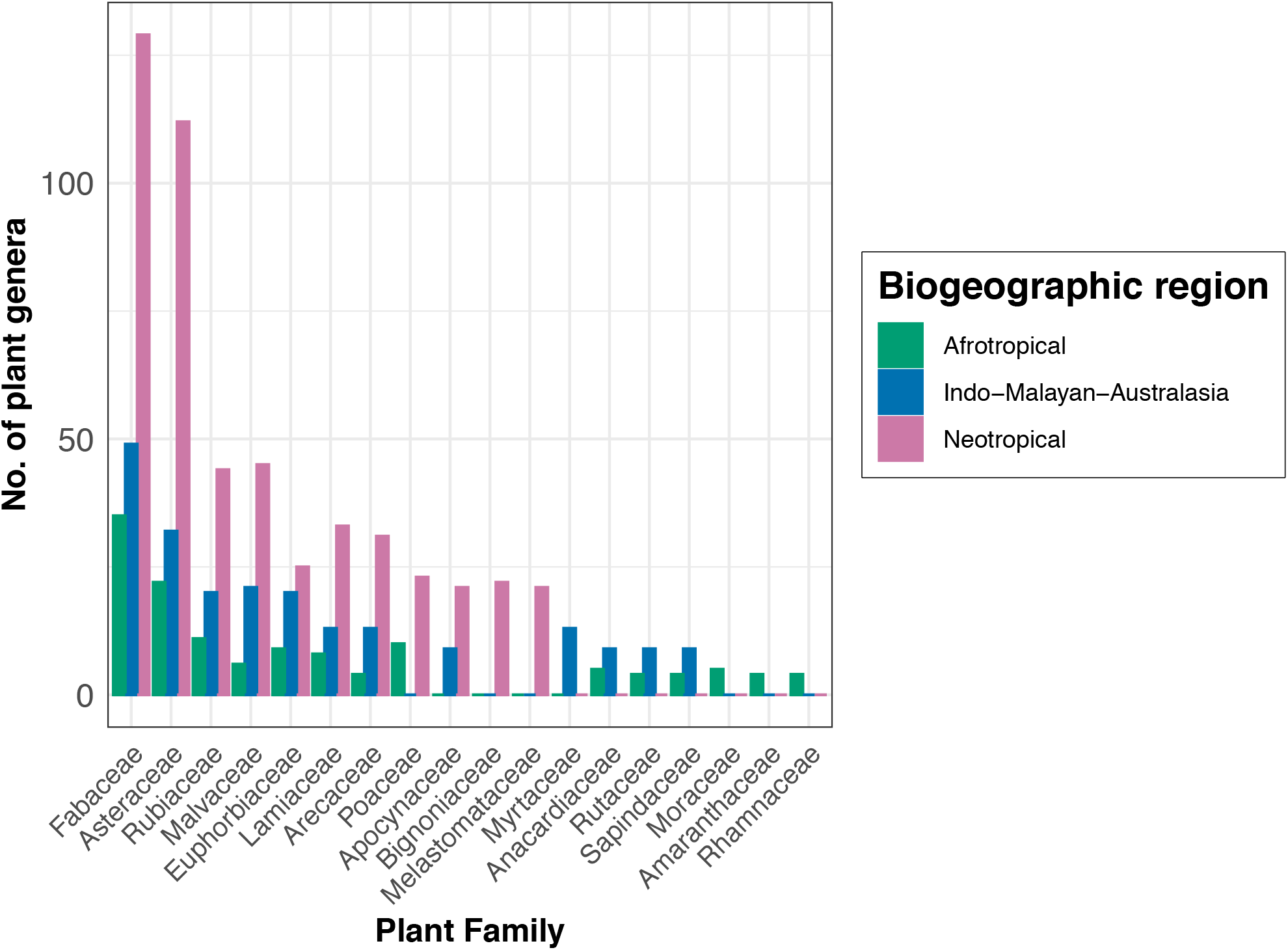
The number of genera visited by stingless bees of the Neotropical, Afrotropical and Indo-Malayan-Australasian regions, in each of the 10 plant families with the highest number of genera visited, according to reported interactions in the literature (>10 families shown where multiple families had the same rank).

In the Neotropics, the three plant families for which the greatest proportion of total genera occurring in the region are visited by stingless bees were Myrtaceae (stingless bees are reported to visit 52% of all genera described for the Neotropics), Arecaceae (43%) and Sapindaceae (42%) (Sup material, **Table S3**).

Several plant families are reported to be commonly used by the stingless bees of all geographic regions (*e.g*., Fabaceae, Asteraceae, Euphorbiaceae); **Figure 3**. Nevertheless, there were differences in the composition of reported plants (at family level) visited by the stingless bees of each biogeographical region (PERMANOVA: *P*= 0.01, R^2^ = 0.08, **Figure 4**, Sup material **Table S4**). This trend was driven particularly by differences between the floral use of Neotropical and Indo-Malayan-Australasian stingless bee fauna, presumably reflecting differences in the plant communities available to bees in these two regions (*P*= 0.03, R2=0.059). Afrotropical stingless bee floral use (at plant family level) did not differ significantly from either other region (Indo-Malayan-Australasian: *P*= 0.14, R2=0.083; Neotropical: *P*= 0.14, R2=0.041).

**Fig 4.**
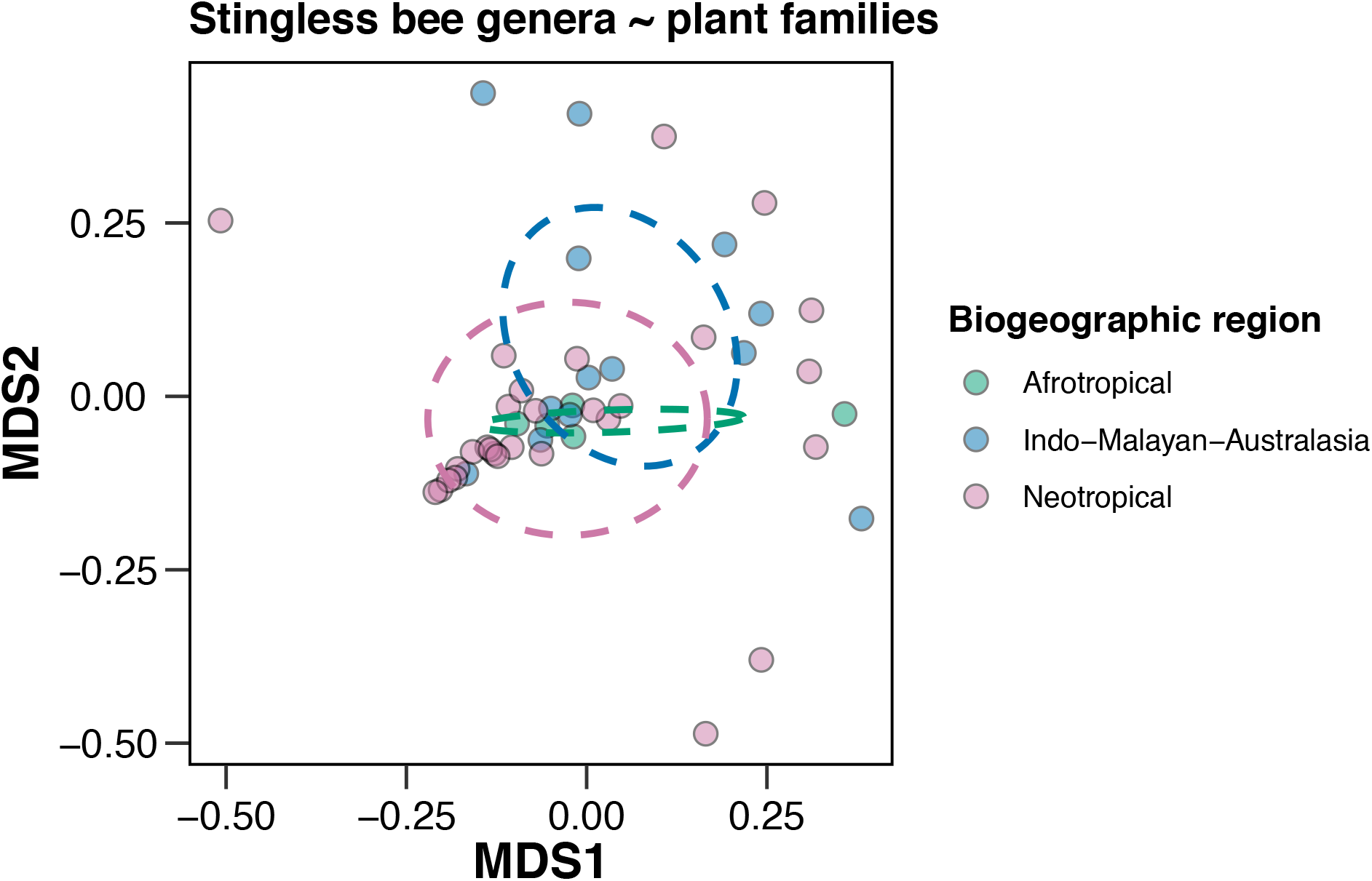
NMDS ordination of stingless bee genera ~ plant family composition in each biogeographic region. Each point represents a bee genera’s floral associations at the plant family level (Afrotropical, N= 5 bee genera; Indo-Malayan-Australasian, N= 13; Neotropical, N= 31). Dashed circles represent the 95% confidence ellipses for each biogeographic region mean (group centroid)

### Traits of highly visited plant genera

7. Across all visited plants, stingless bees visited genera of plants spanning a variety of common types of growth (herbs, trees, shrubs, vines, lianas), including economically important plants such as crops, timber, fibres, medicinal and ornamental use (Sup material, **Table S5**). Within the top 50 plant genera with the most recorded interactions with stingless bees, eight genera of economic importance were visited across all three regions: *Cassia* (Fabaceae), *Citrus* (Rutaceae), *Coffea* (Rubiaceae), *Eucalyptus* (Myrtaceae), *Passiflora* (Passifloraceae), *Psidium* (Myrtaceae), *Senna* (Fabaceae) and *Syzygium* (Myrtaceae).

8. In addition to native flora, stingless bees were reported to visit non-native plants (**Figure 5**). We identified 115 genera of plants that are not native to the region where the bee-plant interaction was reported (57 in the Neotropics, 19 in the Afrotropics and 39 in the Indo-Malayan-Australasian region; Sup material, **Table S6**). Of these 25 (21%) were common edible crops, such as rambutan (*Nephelium* sp.), lychee (*Litchi* sp.) and coffee (*Coffea* sp.) in the Neotropics, mango (*Mangifera* sp.), hog-plum (*Spondias* sp.), and corn (*Zea* sp.) in the Afrotropics, and oil-Palm (*Elaeis* sp.), tamarind (*Tamarindus* sp.) and guava (*Psidium* sp.) in the Indo-Malayan-Australasian region. The remaining genera were non-native garden and ornamental plants, or have been documented as weeds (CABI, 2020); these included numerous genera of Lamiaceae (*e.g*. mint, sage) and Asteraceae (*e.g*. dasies, dandelions, whiteweed).

**Figure 5.**
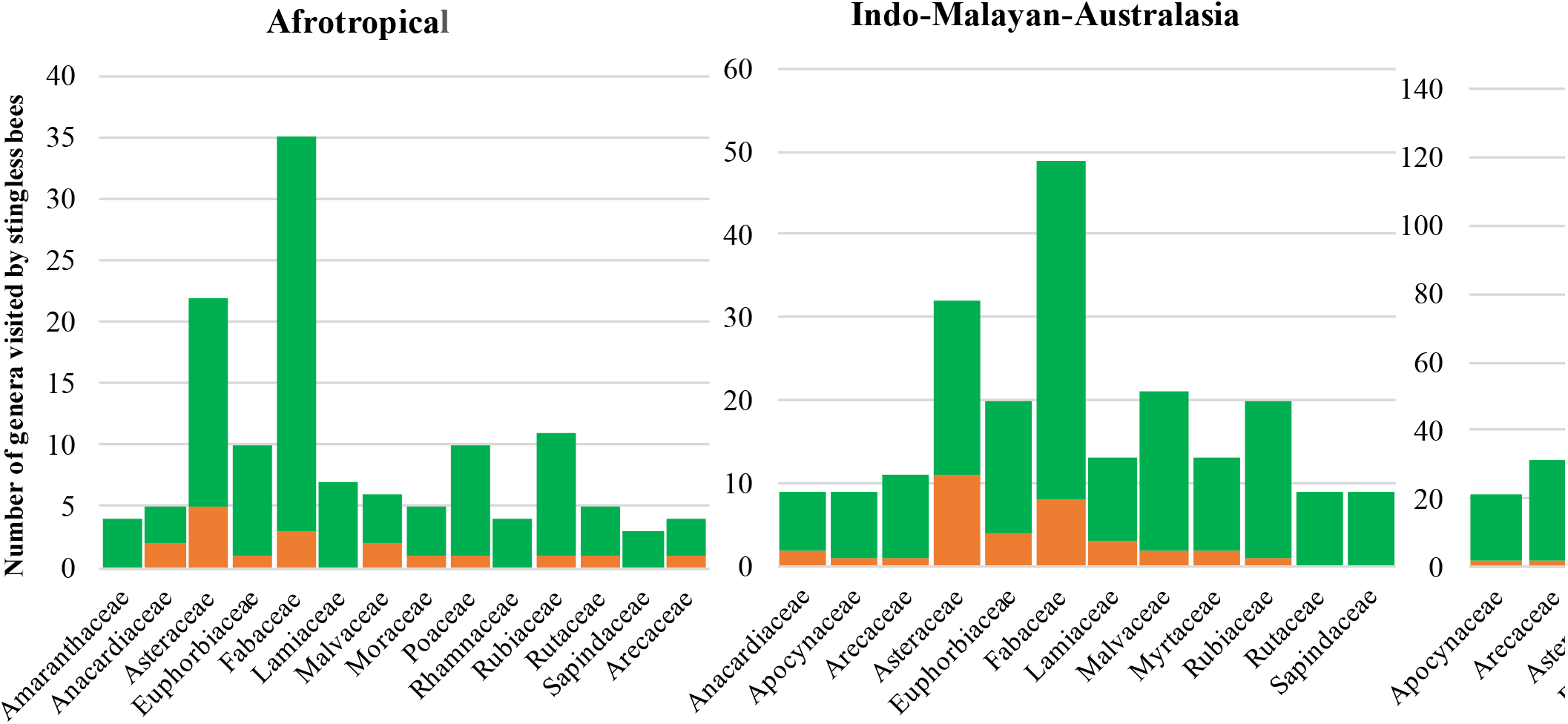
The number of genera visited by stingless bees in the 10 most visited plant families coded by native status, for each of three biogeographical regions: Afrotropical, Indo-Malayan-Australasian and Neotropical (>10 families shown where multiple families had the same rank). Most reported flower visits are for plant genera native to that region (green) but a minority are plant genera introduced to that region (orange).

## Discussion

We consolidated records of floral visitation of wild plants by stingless bees around the world. These records reveal the exceptionally wide variety of plants used as forage by some individual stingless bee species (as many 532 plant genera for *Trigona spinipes*), and by the Meliponini in general. Worldwide, stingless bees visit over 1465 genera from 220 families of flowering plants, around 52% of all angiosperm families (The Plant List, 2010).

This breadth of floral use supports the view that stingless bees play an important role in ecosystem functioning throughout their native range, as pollinators of many tropical and subtropical plants.

### Plants used as forage by stingless bees

We found that the plant families most-used as forage were common to the stingless bee fauna of all three geographic regions: the Neotropics, Afrotropics and Indo-Malayan-Australasian tropics. These included Fabaceae, Asteraceae, Rubiaceae, Malvaceae, Euphorbiaceae, Arecaceae, Lamiaceae and Myrtaceae. The frequent use of several of these plant families has been previously reported for stingless bee species of the Neotropics, including Fabaceae, Asteraceae, Myrtaceae, Malvaceae and Arecaceae (Aleixo et al., 2013; Antonini et al., 2006; Cortopassi-Laurino et al., 1988; Faria et al., 2012; Guibu et al., 1988; Miranda et al., 2015; Ramalho, 1990; Ramalho et al., 1985; Ramalho et al., 1989a). Floral morphology regulates the accessibility to floral resources for animal visitors, and all of the plant families commonly visited by stingless bees have floral traits that have evolved to favour animal visitation, and bee visitation in particular. For instance, the flowers of Myrtaceae, Arecaceae, Asteraceae, Malvaceae and Fabaceae (Subfamily: Mimosoideae) have open corollas with many stamens and longitudinally opened anthers, which facilitates the acquisition of pollen and nectar by bees (Lewis, 2005; Torres et al., 2002). Likewise, animal-pollinated floral resources that are mostly or entirely absent from our database include those with flower types ill-suited to stingless bees. In particular, those with long and narrow corollas (*e.g*., some genera of Lamiaceae) are instead specialized for pollination by one or several species of long-tongued insect visitors (*e.g*., Orchid bees, Tribe Euglossini; (Borrell, 2005; Rodríguez-Gironés et al., 2006)). Stingless bees may still access such resources in some cases by biting holes in corollas, a form of “nectar-robbery” that has been observed in some species (Barrows, 1976). However, more accessible flower forms are likely to be preferred when they are available.

For all three biogeographical regions, Fabaceae and Asteraceae had the most genera visited by stingless bees. As these families are also the two most speciose angiosperm families in the world, and contain many plant species that rely on insect pollination (Christenhusz et al., 2016), their position at the top of the most-used plant families is not surprising. Asteraceae and Fabaceae tend to dominate as bee-forage plants in tropical areas with open vegetation, including where forests have been cleared for human activities (Ramalho et al., 1990). For example, plants in these families are often among the first ones to colonize disturbed or clearer forest areas (Citadini-Zanette et al., 2017; Valdez-Hernández et al., 2014). The ready use of these plants by stingless bees may thus help to explain the resilience of these bees in disturbed habitats, when other sources of forage are not available (Aizen et al., 2012).

While commonly-used families were shared globally, the relative use of different plant families differs between the stingless bees from the Indo-Malayan-Australasian and Neotropical regions. Three possible explanations may account for this result. First, this might reflect biases in the dataset, with a far higher number of reported bee-plant interactions for the Neotropics than other regions. Second, it is likely to indicate, at least in part, differences in the abundance and availability of each plant family in the different regions. For example, bignonias (Bignoniaceae) appear more frequently in visitation records of Neotropical stingless bees than those of other regions, and the greatest diversity of bignonias is also in the Neotropics (Gentry, 1980, 1992). Third, it may reflect genuine differences in the foraging preferences of the stingless bees in each clade, stemming from different coevolutionary histories between the bees and flora of each region.

Foraging preferences in stingless bees can be considered on a variety of scales. At the broadest level, commonly-used plant families across all meliponines can be considered to reflect general preference for those families. The tropics hold a vast range of angiosperms flowering at the same time, making it a diverse and competitive marketplace for the plants. Floral rewards for pollinators differ strongly between plants, and also vary over time (Heinrich, 2004; Willmer et al., 2004), encouraging some level of specialisation even among foragers that use a wide variety of plants. Pollinators thus become receptive to particular flower traits in their search for food, such as flower color, morphology, scent, and temperature (Dyer et al., 2006; Heinrich, 2004; Menzel, 1985). Consistent with this, past studies in the Neotropics have indicated that stingless bees are not indiscriminate generalists, but rather that at least some neotropical genera (*Scaptotrigona* and *Melipona*) show preferences for certain families of plants such as Sapindaceae, Arecaceae, Solanaceae, Myrtaceae, Asteraceae, Melastomataceae Fabaceae and Convolvulaceae (Antonini et al., 2006; da Luz et al., 2019; Ramalho et al., 1989a; Wilms et al., 1997). There is little current information however on how these floral preferences are induced in stingless bees in their natural habitats. As they have perennial nests with overlapping generations, stingless bees may simply favour the most predictable floral resources in their particular locale. For example, in the Neotropics, the most frequently visited species belong to families that flower all year-round (Antonini et al., 2006).

At the level of individual bee species, other factors will also impact foraging preferences. In particular, bee body size is likely to be a key predictor of floral preferences. Body size is related to foraging distance (Araújo et al., 2004; Greenleaf et al., 2007) and determines the spatial scale at which species’ are able to visit flowering plants and tolerate spatial and temporal changes in floral resource availability (Borges et al., 2020). In theory, larger bees might therefore have the scope to be choosier when it comes to floral resources, while smaller species may be more constrained to forage on the plants close to their nest. The Neotropical genera *Melipona* and *Trigona* include some of the largest stingless bees, and these genera also provide some of the clearest evidence for specialisation in floral preferences (Antonini et al., 2006; Nagamitsu et al., 2005; Ramalho et al., 1989b). In a study in Brazil, *Melipona* only visited 21% out of all the plants in flower within their area of foraging. Furthermore, body size and colony size (*i.e*., number of foragers) might also shape floral preferences via competition. For example, species that are small in size, or have few foragers, may choose to exploit flowers less frequently visited by larger, more aggressive bees or colonies (Hubbell et al., 1978; Johnson et al., 1974, 1975; Sommeijer et al., 1983) which tend to monopolize rich resources (Nagamitsu et al., 1999).

### Visitation of non-native plants by stingless bees

Stingless bees are reported to visit many plant genera that are not native to their region, even in areas where native vegetation is preserved (da Luz et al., 2019; Wilson et al., 2021). Stingless bees are thus capable of facilitating the pollination and spread of non-native species (Chytrý et al., 2008; Levine et al., 2004; Marvier et al., 2004). For example, stingless bees in every region visited the genus *Eucalyptus*, which are trees with abundant flowering events that produce high amounts of nectar and pollen, and at any given time of the year one or more species will flower, offering resources practically all year. Eucalypts are native to Australia and cultivated in most tropical regions of the world (Doughty, 2000). Plantations of Eucalypt for timber production in many parts of the world cause habitat fragmentation and loss of biodiversity (Williams, 2015). However they may also provide a safe source of food for bees in degraded ecosystems (Hilgert-Moreira et al., 2014). In addition to Eucalypt, Neotropical stingless bees also forage on at least eight genera of Lamiaceae that are not native to their continent, and the stingless bees of Indo-Malayan-Australasia forage on at least 11 introduced genera of Asteraceae. Many species in these two families are considered invasive weeds. For instance, *Sonchus oleraceus* (Asteraceae) and *Leonorus sibiricus* (Lamiaceae) are catalogued as some of the worst to control weeds due to their quick life cycles and production of highly dispersive seeds (Holm et al., 1977; Kwon et al., 2016; Peerzada et al., 2019).

Yet the willingness of stingless bees to use novel resources, including invasive plants, is also what makes these bees effective pollinators of some crops. Thus, stingless bees in our database are reported to visit and pollinate a diverse range of economically important plants that are not native to their continent. In particular, across all regions they visit economically important crops with mass-blooming phenology, such as coffee (*Coffea*), guava (*Psidium*), mango (*Mangifera*), hog-plum (*Spondias*), tamarind (*Tamarindus*), passion fruit (*Passiflora*) and guava (*Psidium*). The fact that stingless bees across all regions visit these crops highlights potential new opportunities for local pollination management programs using stingless bees globally in these crops.

### Gaps in our knowledge of stingless bee-plant interactions

Our database highlights two key gaps in our knowledge of the interactions between stingless bees and wild plants. First, to what extent do stingless bees contribute to the reproduction of the many plants they visit? While floral visitation often correlates to pollination (Ballantyne et al., 2017; de Santiago-Hernández et al., 2019), not all floral visitation results in effective pollen transfer, and not all pollinators are equally efficient on a per visit basis (King et al., 2013). For example, small-bodied stingless bees that visit large flowers may often fail to contact the flower’s stigma when collecting pollen (Gruter 2020). Pollination studies are time-intensive and difficult in the field, and most confirmed pollination by stingless bees is for crop species (Slaa et al., 2006). Where pollination of wild plants is considered, it is often in regions adjacent to crops. How often stingless bees impact plant reproduction in other ways is also poorly understood. For example, several stingless bees are seed dispersers for plants (mellitochory) which have evolved seeds that offer rewards to the bees in the form of resin (*i.e*., nesting material (Garcia et al., 1992; Wallace et al., 1995)).

Second, our database shows that floral visitation for many of the world’s stingless bees is still undocumented. We identified published records for just half (53%) of all stingless bee species (287 species). Even in Brazil, where decades of research has pioneered knowledge of stingless bee behaviour and ecology (Grüter, 2020), the foraging ecology of much of the country’s rich stingless bee diversity is unknown (Campbell et al., 2019). Data on stingless bee floral use from the Indo-Malayan-Australasian and Afrotropical regions is particularly sparse. With the exception of Australia and New Guinea, honey bees (*Apis* spp) are also native to these regions and are often the focus of traditional beekeeping practices. For example, the Asian hive bee *A. cerana* is widely kept for honey and pollination throughout Asia and India (Oldroyd et al., 2009), while the Western honey bee *A. mellifera* is native throughout Africa. Perhaps for this reason, research interest in stingless bees as pollinators (of both crops and native plants) has lagged behind South and Central America in recent decades. Research of the bees of these regions is now steadily increasing and is certain to advance further in coming years. Some tropical regions in which stingless bee diversity is highest face significant long-term challenges to conserving their ecosystems and pollination services, including high levels of poverty and habitat degradation (Bradshaw et al., 2009). In addition, many stingless bee species in Africa, India, Asia and Australia are cryptic in their morphology, and thus difficult to identify reliably in the field. For example, in Australia, three common species with overlapping distributions are identical in forager appearance, though each are genetically distinct and build unique nest structures (*Tetragonula* sp. of the “carbonaria complex”; (Dollin et al., 1997)).

### Conclusion

Tropical regions support many biodiversity hotspots (Myers et al., 2000), including a high diversity and endemism of both stingless bees and flowering plants (Antonelli et al., 2015; Hawkins et al., 2011). Just as stingless bees rely on plants as food sources for pollen and nectar, plants rely on stingless bees and other pollinators for reproduction. Here we consider stingless bee-plant interactions at a global scale, focusing on patterns at the level of bee genera and plant families. Ultimately, a rich understanding of stingless bee floral use will require continued study at local levels for the many and diverse ecosystems of the tropics and subtropics. Our database aims to provide an easy-reference and helpful initial resource for such future studies on the interactions between wild endemic, endangered or invasive plant species and their stingless bee visitors.

## Acknowledgements

RG is supported by an Australian Research Council Award (DE220100466). The Conselho Nacional de Desenvolvimento Científico e Tecnológico (CNPq, 149154/2018-6 and 380653/2021-4 to DAA).

## Notes

https://datadryad.org/stash/share/XDmAqN_sxG2qf91qiK040SOHlRP4IKrsWsmoX00UxRA.

### Competing Interest Statement

The authors have declared no competing interest.

### Summary of Updates

We have edited some paragraphs and updated the figures and tables.

