## Supplemental tables and figures for "Stingless bee floral visitation in the global tropics and subtropics"

**Supplementary material**

Table S1. Database of reported interactions between stingless bees and flowering plants. This database can be accessed here:

https://datadryad.org/stash/share/XDmAqN_sxG2qf91qiK040SOHlRP4IKrsWsmoX00UxRA

Table S2. The 15 plant families with the most reported genera visited by stingless bees worldwide, and for each of three regions (Neotropical, Afrotropical and Indo-Malayan-Australasian); the total genera reported per family worldwide (The Plant List, 2010), the total visited and the proportion of all genera in the plant family known to be visited by stingless bees based on reported interactions in the literature.

| | | Genera | | |
|---|---|---|---|---|
| Region | Plant Family | Visited | Total | Proportion (%) |
| All regions | Acanthaceae | 21 | 225 | 9 |
| All regions | Anacardiaceae | 21 | 72 | 3 |
| All regions | Apocynaceae | 27 | 402 | 7 |
| All regions | Arecaceae | 41 | 187 | 22 |
| All regions | Asteraceae | 134 | 1765 | 8 |
| All regions | Bignoniaceae | 27 | 85 | 32 |
| All regions | Euphorbiaceae | 41 | 229 | 18 |
| All regions | Fabaceae | 161 | 917 | 18 |
| All regions | Lamiaceae | 43 | 250 | 17 |
| All regions | Malvaceae | 56 | 236 | 24 |
| All regions | Melastomataceae | 27 | 153 | 18 |
| All regions | Myrtaceae | 27 | 144 | 19 |
| All regions | Orchidaceae | 22 | 925 | 2 |
| All regions | Poaceae | 34 | 777 | 4 |
| All regions | Rubiaceae | 65 | 617 | 11 |
| All regions | Sapindaceae | 26 | 124 | 21 |
| Neotropical | Acanthaceae | 17 | 225 | 8 |
| Neotropical | Apocynaceae | 21 | 402 | 5 |
| Neotropical | Arecaceae | 31 | 187 | 17 |
| Neotropical | Asteraceae | 112 | 1765 | 6 |
| Neotropical | Bignoniaceae | 22 | 85 | 26 |
| Neotropical | Euphorbiaceae | 25 | 229 | 11 |
| Neotropical | Fabaceae | 129 | 917 | 14 |
| Neotropical | Lamiaceae | 33 | 250 | 13 |
| Neotropical | Malpighiaceae | 17 | 81 | 21 |
| Neotropical | Malvaceae | 45 | 236 | 19 |
| Neotropical | Melastomataceae | 21 | 153 | 14 |
| Neotropical | Myrtaceae | 20 | 144 | 14 |
| Neotropical | Poaceae | 23 | 777 | 3 |
| Neotropical | Rubiaceae | 44 | 617 | 7 |

| | | | | |
|---|---|---|---|---|
| Neotropical | Sapindaceae | 20 | 124 | 16 |
| Indo-Malayan-Australasian | Acanthaceae | 7 | 225 | 3 |
| Indo-Malayan-Australasian | Anacardiaceae | 9 | 72 | 13 |
| Indo-Malayan-Australasian | Apocynaceae | 9 | 402 | 2 |
| Indo-Malayan-Australasian | Arecaceae | 13 | 187 | 7 |
| Indo-Malayan-Australasian | Asteraceae | 32 | 1765 | 2 |
| Indo-Malayan-Australasian | Cucurbitaceae | 8 | 126 | 6 |
| Indo-Malayan-Australasian | Euphorbiaceae | 20 | 229 | 9 |
| Indo-Malayan-Australasian | Fabaceae | 49 | 917 | 5 |
| Indo-Malayan-Australasian | Lamiaceae | 13 | 250 | 5 |
| Indo-Malayan-Australasian | Malvaceae | 21 | 236 | 9 |
| Indo-Malayan-Australasian | Myrtaceae | 13 | 144 | 9 |
| Indo-Malayan-Australasian | Orchidaceae | 8 | 925 | 1 |
| Indo-Malayan-Australasian | Rubiaceae | 20 | 617 | 3 |
| Indo-Malayan-Australasian | Rutaceae | 9 | 148 | 6 |
| Indo-Malayan-Australasian | Sapindaceae | 9 | 124 | 7 |
| Afrotropical | Acanthaceae | 3 | 225 | 1 |
| Afrotropical | Amaranthaceae | 4 | 163 | 2 |
| Afrotropical | Amaryllidaceae | 3 | 79 | 4 |
| Afrotropical | Anacardiaceae | 5 | 72 | 7 |
| Afrotropical | Arecaceae | 4 | 187 | 2 |
| Afrotropical | Asteraceae | 22 | 1765 | 1 |
| Afrotropical | Connaraceae | 3 | 16 | 19 |
| Afrotropical | Cucurbitaceae | 3 | 126 | 2 |
| Afrotropical | Euphorbiaceae | 9 | 229 | 4 |
| Afrotropical | Fabaceae | 35 | 917 | 4 |
| Afrotropical | Lamiaceae | 8 | 250 | 3 |
| Afrotropical | Malvaceae | 6 | 236 | 3 |
| Afrotropical | Moraceae | 5 | 37 | 14 |
| Afrotropical | Myrtaceae | 3 | 144 | 2 |
| Afrotropical | Orchidaceae | 3 | 925 | 1 |
| Afrotropical | Phyllanthaceae | 3 | 58 | 5 |
| Afrotropical | Poaceae | 10 | 777 | 1 |
| Afrotropical | Rhamnaceae | 4 | 52 | 8 |
| Afrotropical | Rubiaceae | 11 | 617 | 2 |
| Afrotropical | Rutaceae | 4 | 148 | 3 |
| Afrotropical | Sapindaceae | 4 | 124 | 3 |
| Afrotropical | Solanaceae | 3 | 105 | 3 |
| Afrotropical | Verbenaceae | 3 | 34 | 9 |

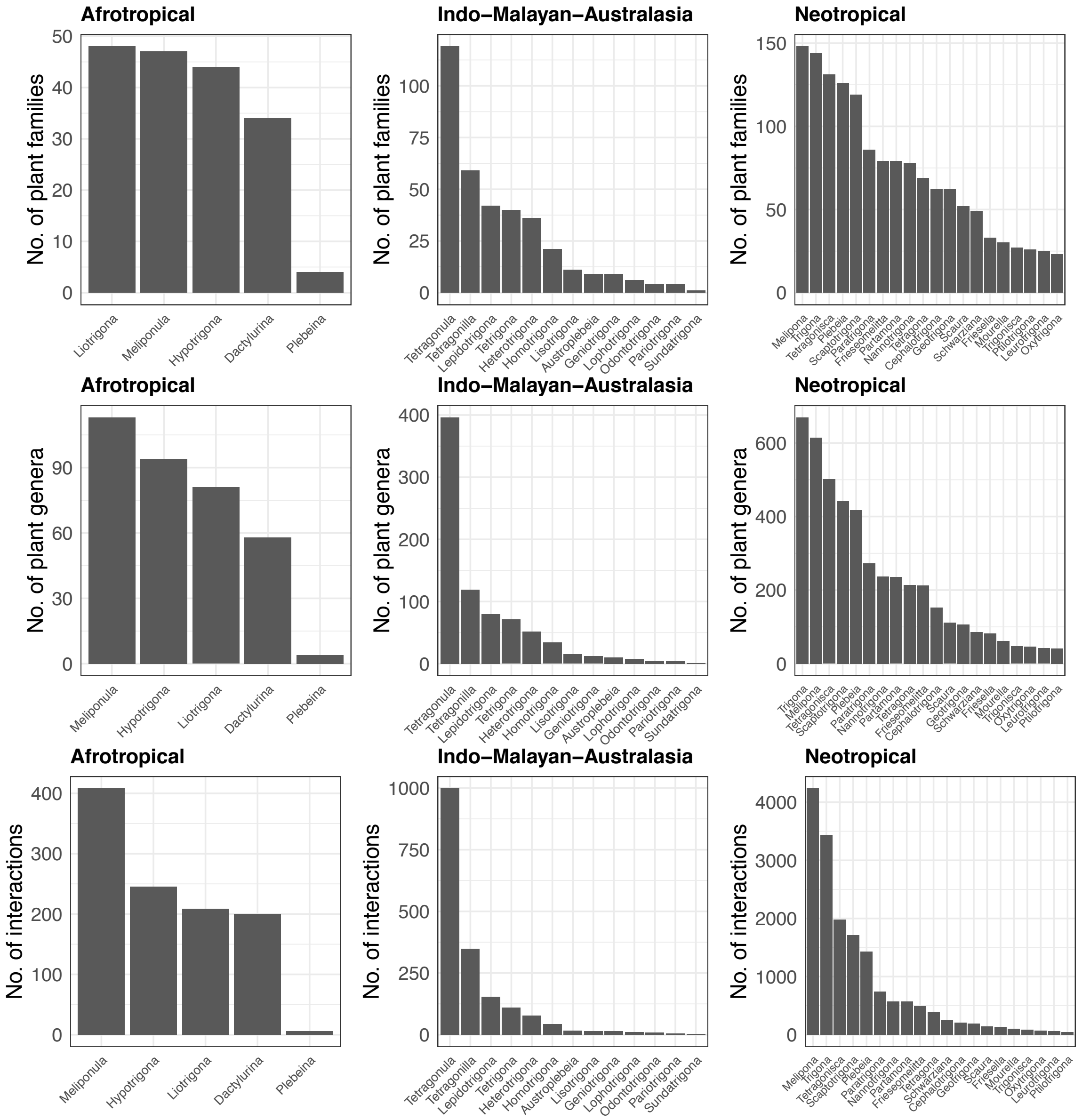

Fig. S1. Histograms with the counts a of plant families visited by stingless bee genera per geographical region. b) of plant genera visited by stingless bee genera per geographical region c) with the total number of plant and stingless bee interactions ordered by stingless bee genera.

Table S3. The 15 plant families with the most reported genera visited by stingless bees in the Neotropics; and the proportion of all genera reported for the Neotropics (Milliken, 2009) in the plant family known to be visited by stingless bees, based on reported interactions in the literature. The * shows the three families with the highest proportion of genera visited in the Neotropics.

| | Genera | | Genera visited in the Neotropics | | |
|---|---|---|---|---|---|
| Plant Family | Worldwide | Neotropics | Native | Non-native | Proportion (%) |
| Acanthaceae | 225 | NA | 15 | 2 | NA |
| Apocynaceae | 402 | 50 | 19 | 2 | 38 |
| Arecaceae | 187 | 68 | 29 | 2 | 43* |
| Asteraceae | 1765 | NA | 100 | 12 | 17 |
| Bignoniaceae | 85 | 77 | 20 | 2 | 26 |
| Euphorbiaceae | 229 | 82 | 23 | 2 | 28 |
| Fabaceae | 917 | 314 | 120 | 8 | 38 |
| Lamiaceae | 250 | 65 | 25 | 8 | 38 |
| Malpighiaceae | 81 | 59 | 17 | 0 | 29 |
| Malvaceae | 236 | 129 | 41 | 4 | 32 |
| Melastomataceae | 153 | 107 | 21 | 0 | 20 |
| Myrtaceae | 144 | 29 | 15 | 5 | 52* |
| Poaceae | 777 | 288 | 20 | 3 | 7 |
| Rubiaceae | 617 | 220 | 41 | 3 | 19 |
| Sapindaceae | 124 | 38 | 16 | 4 | 42* |

Table S4. PERMANOVA (Permutational Multivariate Analysis of Variance) results comparing the differences in floral preferences (plant family) between the three biogeographical subclades of stingless bees (Afrotropical, Indo-Malayan-Australasian and Neotropics), based on reported bee-plant interactions in the literature.

| | F.Model | $R^2$ | *P* (adjusted) |
|---|---|---|---|
| Afrotropical vs Indo-Malayan-Australasia | 1.44783591 | 0.08298083 | 0.144 |
| Afrotropical vs Neotropical | 1.396818549 | 0.041824899 | 0.144 |
| Indo-Malayan-Australasia vs Neotropical | 2.519527321 | 0.05925577 | 0.03* |

Table S5. The top 50 Plant genera with the most recorded visitations by stingless bees in three biogeographical regions (POWO, 2019). * IMAA region = Indo-Malayan-Australasian

| Region* | Family | Genera | Distribution | Growth type | Crop name | Other use |
|---|---|---|---|---|---|---|
| Afrotropical/IMAA | Fabaceae | Acacia | Native | Shrub/Tree | | Timber and sap |
| IMAA | Asteraceae | Acmella | Native | Herb | | |
| Afrotropical | Passifloraceae | Adenia | Native | Herb/Shrub/Tree/ Liana/Vine | | |
| Afrotropical/IMAA | Asteraceae | Ageratum | Non-native | Herb/Shrub | | Ornamental and medicinal |
| Neotropical | Euphorbiaceae | Alchornea | Native | Shrub/Tree | | |
| IMAA | Amaryllidaceae | Allium | Native | Herb | Onion, garlic, scallion, shallot, leek, and chives. | Medicinal |
| Afrotropical | Amaranthaceae | Amaranthus | Native | Herb | Grains | |
| Afrotropical | Annonaceae | Annona | Native | | Custard apple, soursop | Medicinal |
| IMAA | Phyllanthaceae | Antidesma | Native | Shrub/Tree | | |
| Afrotropical | Polygonaceae | Antigonon | Non-native | Vines | | |
| IMAA | Primulaceae | Ardisia | Native | Shrub/Tree | Drupe fruits | Medicinal |
| Afrotropical/IMAA | Acanthaceae | Asystasia | Native | Herb | | Ornamental and food |
| Afrotropical/IMAA | Oxalidaceae | Averrhoa | Non-Native | Shrub/Tree | Star-fruit | |
| IMAA | Meliaceae | Azadirachta | Native | Tree | Neem oil | Medicinal |
| Neotropical | Asteraceae | Baccharis | Native | Shrub | | |
| Afrotropical/Neotropical | Fabaceae | Bauhinia | Native | Tree | | Ornamental |
| Afrotropical/IMAA | Asteraceae | Bidens | Native | Herb | | |
| Neotropical | Bixaceae | Bixa | Native | Shrub | Annatto | |
| IMAA | Euphorbiaceae | Blumeodendron | Native | Tree | | |
| Afrotropical | Arecaceae | Borassus | Native | Tree | | Leaves for crafts and sweet sap |
| IMAA | Brassicaceae | Brassica | Native | Herb/Shrub | Canola, brown mustard, turnip, cabbage, cauliflower, etc. | |
| Afrotropical | Asteraceae | Brenandendron | Native | Herb | | |

| | | | | | | |
|---|---|---|---|---|---|---|
| Afrotropical | Phyllanthaceae | Bridelia | Native | Shrub/Tree | | Medicinal |
| Neotropical | Malpighiaceae | Byrsonima | Native | Shrub/Tree | Nance | |
| Afrotropical/Neotropical | Fabaceae | Caesalpinia | Native | Tree/Shrub/Liana | | Ornamental |
| Afrotropical | Fabaceae | Calliandra | Native | Shrub/Tree | | Ornamental |
| IMAA | Convolvulaceae | Camonea | Native | Herb | | Ornamental |
| IMAA/Neotropical | Solanaceae | Capsicum | Non-native | Herb/Shrub | Pepper and Chilli | |
| Afrotropical | Rubiaceae | Casasia | Non-native | Shrub/Tree | Edible fruits | Timber |
| Neotropical | Salicaceae | Casearia | Native | Shrub/Tree | Chilli | Medicinal |
| Afrotropical/IMAA/Neo | Fabaceae | Cassia | Native | Tree/Shrub | | Ornamental and use in reforestation projects |
| Neotropical | Urticaceae | Cecropia | Native | Tree | | Ornamental and use in reforestation projects |
| Neotropical | Fabaceae | Chamaecrista | Native | Tree/Shrub | | Ornamental and use in reforestation projects |
| Afrotropical | Vitaceae | Cissus | Native | Liana | | Ornamental/Medicinal |
| Afrotropical/IMAA/Neo | Rutaceae | Citrus | Non-native | Tree/Shrub | Citrus fruits | Ornamental |
| Neotropical | Polygonaceae | Coccoloba | Native | Shrub/Tree/Liana | | |
| Afrotropical | Bixaceae | Cochlospermum | Native | Shrub/Tree | | |
| IMAA/Neotropical | Arecaceae | Cocos | Native | Tree | Coconut | Ornamental |
| Afrotropical/IMAA/Neo | Rubiaceae | Coffea | Native | Shrub | Coffee | |
| Afrotropical | Lamiaceae | Coleus | Native | Herb/Shrub | | Ornamental |
| Afrotropical | Combretaceae | Combretum | Native | Shrub/Tree | | Medicinal |
| Neotropical | Boraginaceae | Cordia | Native | Shrub/Tree | | Medicinal, Timber and sap |
| Afrotropical | Rubiaceae | Crossopteryx | Native | Shrub/Tree | | Medicinal |
| Afrotropical/IMAA | Fabaceae | Crotalaria | Native | Herb/Shrub | Mitoo | Ornamental |
| IMAA/Neotropical | Euphorbiaceae | Croton | Native | Herb/Shrub/Tree/Liana | | Ornamental and use of bark |
| Neotropical | Cucurbitaceae | Cucurbita | Native | Herb/Vine | Squash, pumpkin, cucumber, etc. | |
| Neotropical | Sapindaceae | Cupania | Native | Shrub/Tree | | |
| Afrotropical | Araliaceae | Cussonia | Native | Shrub/Tree | | Medicinal |
| Afrotropical | Cyperaceae | Cyperus | Native | Herb | | |
| IMAA | Orchidaceae | Dendrobium | Native | Herb | | Ornamental |

| | | | | | | |
|---|---|---|---|---|---|---|
| Afrotropical | Euphorbiaceae | Dichostemma | Native | Shrub/Tree | | |
| IMAA | Dilleniaceae | Dillenia | Native | Shrub/Tree | | |
| IMAA | Ebenaceae | Diospyros | Native | Shrub/Tree | Persimmon | Ornamental and use of bark |
| Neotropical | Verbenaceae | Duranta | Native | Shrub/Tree | | Ornamental |
| Afrotropical | Asteraceae | Elephantopus | Native | Herb | | Medicinal |
| Afrotropical | Fabaceae | Entada | Native | Shrub/Tree/Liana | | Medicinal |
| Afrotropical/IMAA/Neo | Myrtaceae | Eucalyptus | Non-native | Shrub/Tree | | Ornamental and medicinal |
| IMAA/Neotropical | Myrtaceae | Eugenia | Native | Shrub/Tree | Edible fruit | Ornamental |
| Afrotropical/Neotropical | Euphorbiaceae | Euphorbia | Native | Herb/Shrub/Tree | | Ornamental |
| Neotropical | Arecaceae | Euterpe | Native | Tree | Açai berry | |
| IMAA | Moraceae | Ficus | Native | Tree/Shrub/Vine | | Ornamental |
| Neotropical | Rosaceae | Fragaria | Native | Herb | Strawberry | |
| Afrotropical | Rhamnaceae | Gouania | Native | Shrub/Liana | | |
| IMAA | Tiliaceae | Grewia | Native | Shrub/Tree | | Medicinal |
| Afrotropical | Asteraceae | Guizotia | Native | Herb | Oil and edible seeds | |
| Afrotropical | Hypericaceae) | Harungana | Native | Shrub/Tree | | Medicinal |
| IMAA/Neotropical | Asteraceae | Helianthus | Non-native | Herb | Edible seeds and oil | Ornamental |
| IMAA | Malvaceae | Hibiscus | Native | Herb/Shrub/Tree | Edible flowers | Ornamental and fibres |
| Afrotropical | Fabaceae | Indigofera | Native | Herb/Shrub/Tree | | Ornamental |
| Neotropical | Fabaceae | Inga | Native | Shrub/Tree | | Ornamental |
| Afrotropical/IMAA | Convolvulaceae | Ipomoea | Native | Herb/Shrub/Tree/Liana | | Ornamental, sweet potato and water spinach |
| IMAA | Rubiaceae | Ixora | Native | Shrub/Tree | | Ornamental |
| IMAA | Euphorbiaceae | Jatropha | Native | Shrub/Tree | | Fibres |
| IMAA | Lythraceae | Lagerstroemia | Native | Shrub/Tree | | Ornamental |
| Afrotropical/IMAA | Fabaceae | Leucaena | Non-native | Tree/Shrub | Edible fruits | Timber and reforestation |
| Afrotropical/IMAA | Anacardiaceae | Mangifera | Non-native | Shrub/Tree | Mango | |
| Afrotropical | Bignoniaceae | Markhamia | Native | Shrub/Tree | | Medicinal |
| IMAA | Myrtaceae | Melaleuca | Native | Shrub/Tree | | Ornamental and oil |

| | | | | | | |
|---|---|---|---|---|---|---|
| IMAA | Melastomataceae | Melastoma | Native | Shrub | | Ornamental |
| Neotropical | Melastomataceae | Miconia | Native | Shrub/Tree | | Timber |
| Neotropical | Asteraceae | Mikania | Native | Herb | | Medicinal |
| IMAA | Fabaceae | Millettia | Native | Shrub/Tree | | |
| IMAA/Neotropical | Fabaceae | Mimosa | Native | Herb/Shrub | | Ornamental, pioneer trees and Timber |
| IMAA | Muntingiaceae | Muntingia | Non-native | Shrub/Tree | Edible fruit | Medicinal |
| IMAA | Commelineae | Murdannia | Native | Herb | | |
| Afrotropical | Musaceae | Musa | Non-native | Herb | Banana | |
| Neotropical | Myrtaceae | Myrcia | Native | Shrub/Tree | | |
| Afrotropical | Rubiaceae | Nauclea | Native | Shrub/Tree | | Timber |
| IMAA | Sapindaceae | Nephelium | Native | Shrub/Tree | Litchi | |
| Neotropical | Lauraceae | Ocotea | Native | | | Timber |
| Afrotropical | Burseraceae | Pachylobus | Native | Shrub/Tree | African plum | |
| Afrotropical/IMAA/Neo | Passifloraceae | Passiflora | Native | Herb/Shrub/Vine | Passion fruit | Ornamental |
| IMAA | Fabaceae | Peltophorum | Native | Shrub/Tree | | Medicinal and Timber |
| Neotropical | Lauraceae | Persea | Native | Shrub/Tree | Avocado | |
| Afrotropical | Polygonaceae | Persicaria | Native | Herb | | Ornamental |
| IMAA | Urticaceae | Pipturus | Native | Shrub/Tree | | Medicinal |
| Neotropical | Melastomataceae | Pleroma | Native | Shrub/Tree | | Ornamental |
| Neotropical | Burseraceae | Protium | Native | Shrub/Tree | Edible fruit | Medicinal and Timber |
| IMAA | Rosaceae | Prunus | Native | | Apricot, plum, peach, etc. | Ornamental and Timber |
| Afrotropical/IMAA/Neo | Myrtaceae | Psidium | Non-native | Tree/Shrub | Guava | |
| IMAA | Rosaceae | Rubus | Native | Herb/Shrub | Raspberry | |
| IMAA | Burseraceae | Santiria | Native | Shrub/Tree | | |
| Neotropical | Araliaceae | Schefflera | Native | Shrub/Tree | | |
| Neotropical | Anacardiaceae | Schinus | Native | Shrub/Tree | | Medicinal |
| Afrotropical/IMAA/Neo | Fabaceae | Senna | Native | Herb/Shrub/Tree | | Ornamental |
| Neotropical | Sapindaceae | Serjania | Native | Liana/Vine | | |

| | | | | | | |
|---|---|---|---|---|---|---|
| Afrotropical | Pedaliaceae | Sesamum | Native | Herb | Sesame seeds | |
| Neotropical | Malvaceae | Sida | Native | Herb/Shrub/Tree | | |
| IMAA/Neotropical | Solanaceae | Solanum | Native | Herb/Tree/Shrub/ Vine | Tomato, eggplant, potato | Ornamental |
| Afrotropical/Neotropical | Rubiaceae | Spermacoce | Native | Herb | | Medicinal |
| Neotropical | Anacardiaceae | Spondias | Native | Shrub/Tree | Hog plum | |
| Afrotropical | Verbenaceae | Stachytarpheta | Native | Herb/Tree/Shrub/Vine | | |
| Neotropical | Loranthaceae | Struthanthus | Native | Shrub | | Medicinal |
| Neotropical | Arecaceae | Syagrus | Native | Tree | Edible fruit | |
| IMAA | Asteraceae | Synotis | Native | Herb | | |
| Afrotropical/IMAA/Neo | Myrtaceae | Syzygium | Native | Shrub/Tree | Roseapple | |
| IMAA | Lamiaceae | Tectona | Native | Shrub/Tree | | Teak timber |
| IMAA | Combretaceae | Terminalia | Native | Shrub/Tree | | Medicinal |
| Neotropical | Malvaceae | Theobroma | Native | Shrub/Tree | Cacao | |
| IMAA | Acanthaceae | Thunbergia | Native | Herb/Shrub | | |
| IMAA | Asteraceae | Tridax | Non-native | Herb | | Medicinal |
| Afrotropical | Malvaceae | Urena | Native | Shrub/Tree | | Medicinal |
| Afrotropical | Fabaceae | Vachellia | Native | Shrub/Tree | | Medicinal |
| Afrotropical/Neotropical | Asteraceae | Vernonia | Native | Herb/Shrub/Tree/Liana | | Medicinal, edible leaves, ornamental |
| IMAA | Sapindaceae | Xerospermum | Native | Shrub/Tree | | |
| Afrotropical | Poaceae | Zea | Non-native | Herb | Corn | |

Table S6. List of invasive plant genera (POWO, 2019) from the 15 families with the most visited genera. visited by the stingless bees from the three biogeographical regions. The common edible crop names are at the top of each region.

| Region | Family | Non-native genera | Crop name |
|---|---|---|---|
| Region | Family | *Non-native genera* | Crop name |
| Afrotropical | Anacardiaceae | *Mangifera* | Mango |
| Afrotropical | Anacardiaceae | *Spondias* | Hog-plum |
| Afrotropical | Arecaceae | *Cocos* | Coconut |
| Afrotropical | Asteraceae | *Ageratum* | |
| Afrotropical | Asteraceae | *Flaveria* | |
| Afrotropical | Asteraceae | *Galinsoga* | |
| Afrotropical | Asteraceae | *Synedrella* | |
| Afrotropical | Asteraceae | *Vernonanthura* | |
| Afrotropical | Euphorbiaceae | *Manihot* | |
| Afrotropical | Fabaceae | *Caesalpinia* | |
| Afrotropical | Fabaceae | *Calliandra* | |
| Afrotropical | Fabaceae | *Leucaena* | |
| Afrotropical | Malvaceae | *Ceiba* | |
| Afrotropical | Malvaceae | *Pachira* | |
| Afrotropical | Moraceae | *Artocarpus* | Jackfruit |
| Afrotropical | Myrtaceae | *Psidium* | Guava |
| Afrotropical | Poaceae | *Zea* | Corn |
| Afrotropical | Rubiaceae | *Casasia* | |
| Afrotropical | Rutaceae | *Citrus* | Citrus fruit |
| Indo-Malayan-Australasian | Amaranthaceae | *Spinacia* | |
| Indo-Malayan-Australasian | Amaryllidaceae | *Hippeastrum* | |
| Indo-Malayan-Australasian | Amaryllidaceae | *Nerine* | |
| Indo-Malayan-Australasian | Anacardiaceae | *Schinus* | |
| Indo-Malayan-Australasian | Anacardiaceae | *Tapirira* | Tamarind |
| Indo-Malayan-Australasian | Apocynaceae | *Thevetia* | |
| Indo-Malayan-Australasian | Arecaceae | *Elaeis* | Oil-Palm |
| Indo-Malayan-Australasian | Asteraceae | *Ageratum* | |
| Indo-Malayan-Australasian | Asteraceae | *Calendula* | |
| Indo-Malayan-Australasian | Asteraceae | *Chromolaena* | |
| Indo-Malayan-Australasian | Asteraceae | *Cosmos* | |
| Indo-Malayan-Australasian | Asteraceae | *Crassocephalum* | |
| Indo-Malayan-Australasian | Asteraceae | *Gaillardia* | |
| Indo-Malayan-Australasian | Asteraceae | *Guizotia* | |
| Indo-Malayan-Australasian | Asteraceae | *Helianthus* | Sunflower seeds |
| Indo-Malayan-Australasian | Asteraceae | *Tagetes* | |
| Indo-Malayan-Australasian | Asteraceae | *Tridax* | |
| Indo-Malayan-Australasian | Asteraceae | *Verbesina* | |

| | | | |
|---|---|---|---|
| Indo-Malayan-Australasian | Cucurbitaceae | *Cucurbita* | Cucurbits |
| Indo-Malayan-Australasian | Euphorbiaceae | *Hevea* | |
| Indo-Malayan-Australasian | Euphorbiaceae | *Manihot* | |
| Indo-Malayan-Australasian | Euphorbiaceae | *Ricinus* | |
| Indo-Malayan-Australasian | Euphorbiaceae | *Sapium* | |
| Indo-Malayan-Australasian | Fabaceae | *Caesalpinia* | |
| Indo-Malayan-Australasian | Fabaceae | *Calliandra* | |
| Indo-Malayan-Australasian | Fabaceae | *Delonix* | |
| Indo-Malayan-Australasian | Fabaceae | *Gliricidia* | |
| Indo-Malayan-Australasian | Fabaceae | *Leucaena* | |
| Indo-Malayan-Australasian | Fabaceae | *Phaseolus* | |
| Indo-Malayan-Australasian | Fabaceae | *Psophocarpus* | Guava |
| Indo-Malayan-Australasian | Fabaceae | *Tamarindus* | |
| Indo-Malayan-Australasian | Lamiaceae | *Anisomeles* | |
| Indo-Malayan-Australasian | Lamiaceae | *Clerodendrum* | |
| Indo-Malayan-Australasian | Lamiaceae | *Mesosphaerum* | |
| Indo-Malayan-Australasian | Malvaceae | *Ceiba* | |
| Indo-Malayan-Australasian | Malvaceae | *Malvastrum* | |
| Indo-Malayan-Australasian | Myrtaceae | *Psidium* | Bean |
| Indo-Malayan-Australasian | Myrtaceae | *Syncarpia* | Spinach |
| Indo-Malayan-Australasian | Rubiaceae | *Hamelia* | |
| Neotropical | Acanthaceae | *Geissomeria* | |
| Neotropical | Acanthaceae | *Thunbergia* | |
| Neotropical | Apocynaceae | *Catharanthus* | |
| Neotropical | Apocynaceae | *Nerium* | |
| Neotropical | Arecaceae | *Archontophoenix* | |
| Neotropical | Asteraceae | *Bellis* | |
| Neotropical | Asteraceae | *Calendula* | |
| Neotropical | Asteraceae | *Chrysanthemum* | |
| Neotropical | Asteraceae | *Cichorium* | |
| Neotropical | Asteraceae | *Coleostephus* | |
| Neotropical | Asteraceae | *Cyanthillium* | |
| Neotropical | Asteraceae | *Cynara* | |
| Neotropical | Asteraceae | *Gazania* | |
| Neotropical | Asteraceae | *Gerbera* | |
| Neotropical | Asteraceae | *Guizotia* | |
| Neotropical | Asteraceae | *Helianthus* | Sunflower seeds |
| Neotropical | Asteraceae | *Phoenix* | Date |
| Neotropical | Asteraceae | *Sonchus* | |
| Neotropical | Bignoniaceae | *Podranea* | |
| Neotropical | Bignoniaceae | *Spathodea* | |
| Neotropical | Euphorbiaceae | *Triadica* | |
| Neotropical | Euphorbiaceae | *Vernicia* | Tung oil |

| | | | |
|---|---|---|---|
| Neotropical | Fabaceae | *Adenanthera* | |
| Neotropical | Fabaceae | *Alysicarpus* | |
| Neotropical | Fabaceae | *Julbernardia* | |
| Neotropical | Fabaceae | *Lablab* | Hyacinth bean |
| Neotropical | Fabaceae | *Melilotus* | |
| Neotropical | Fabaceae | *Pueraria* | |
| Neotropical | Fabaceae | *Securigera* | |
| Neotropical | Fabaceae | *Spartium* | |
| Neotropical | Lamiaceae | *Coleus* | |
| Neotropical | Lamiaceae | *Congea* | |
| Neotropical | Lamiaceae | *Holmskioldia* | |
| Neotropical | Lamiaceae | *Leonurus* | |
| Neotropical | Lamiaceae | *Melissa* | |
| Neotropical | Lamiaceae | *Origanum* | Oregano |
| Neotropical | Lamiaceae | *Tectona* | |
| Neotropical | Lamiaceae | *Tetradenia* | |
| Neotropical | Malvaceae | *Abelmoschus* | Okra |
| Neotropical | Malvaceae | *Bombax* | |
| Neotropical | Malvaceae | *Dombeya* | |
| Neotropical | Malvaceae | *Grewia* | |
| Neotropical | Myrtaceae | *Callistemon* | |
| Neotropical | Myrtaceae | *Corymbia* | |
| Neotropical | Myrtaceae | *Eucalyptus* | |
| Neotropical | Myrtaceae | *Melaleuca* | Tea-tree oil |
| Neotropical | Myrtaceae | *Syzygium* | |
| Neotropical | Poaceae | *Coix* | Job's tears grain |
| Neotropical | Poaceae | *Schizostachyum* | |
| Neotropical | Poeaceae | *Melinis* | |
| Neotropical | Rubiaceae | *Coffea* | Coffee |
| Neotropical | Rubiaceae | *Gardenia* | |
| Neotropical | Rubiaceae | *Pentas* | |
| Neotropical | Sapindaceae | *Harpullia* | |
| Neotropical | Sapindaceae | *Koelreuteria* | |
| Neotropical | Sapindaceae | *Litchi* | Litchi |
| Neotropical | Sapindaceae | *Nephelium* | Rambutan |
